# Oscillatory Viscoelastic Microfluidics for Efficient Focusing and Separation of Nanoscale Species

**DOI:** 10.1101/668301

**Authors:** Mohammad Asghari, Xiaobao Cao, Bogdan Mateescu, Daniel van Leeuwen, Stavros Stavrakis, Andrew J. deMello

## Abstract

The ability to precisely control particle migration within microfluidic systems is essential for focusing, separating, counting and detecting a wide range of biological species. To date, viscoelastic microfluidic systems have primarily been applied to the focusing, separation and isolation of micron-sized species, with their use in nanoparticle manipulations being underdeveloped and underexplored, due to issues related to nanoparticle diffusivity and a need for extended channel lengths. To overcome such issues, we herein present sheathless oscillatory viscoelastic microfluidics as a method for focusing and separating both micron and sub-micron species. To highlight the efficacy of our approach, we segment our study into three size regimes, namely micron (where characteristic particle dimensions are above 1 μm), sub-micron (where characteristic dimensions are between 1 μm and 100 nm) and nano (where characteristic dimensions are below 100 nm) regimes. Based on the ability to successfully manipulate particles in all these regimes, we demonstrate the successful isolation of p-bodies from biofluids (in the micron regime), the focusing of λ-DNA (in the sub-micron regime) and the focusing of extracellular vesicles (in the nano-regime). Finally, we characterize the physics underlying viscoelastic microflows using a dimensionless number that relates the lateral velocity (due to elastic effects) to the diffusion constant of the species within the viscoelastic carrier fluid. Based on the ability to precisely manipulate species in all three regimes, we expect that sheathless oscillatory viscoelastic microfluidics will provide for significant new opportunities in a range of biological and life science applications.

## Introduction

In recent years, much attention has focused on the characterization of circulating particles within biofluids (1), with a view to improving the sensitivity and specificity of biomarker detection strategies (2,3). For example, small extracellular vesicles (sEVs) have diameters between 80 and 300 nm and are known to circulate within a variety of biofluids. sEVs exhibit protein, DNA and RNA profiles that are modulated during cancer progression and therapy, and thus can be used for non-invasive monitoring of disease progression based on their molecular profile (1,4,5). Unsurprisingly, a number of techniques have been developed to selectively isolate such species from other circulating debris and cells within biofluids. These include methods based on differential centrifugation (6,7), density gradient separation (8), affinity capture (9), size-exclusion chromatography (10) and ultra-filtration (11) or a combination of these (12), which fractionate particle diversity on the basis of size, density and surface markers. However, it is important to note that these approaches are cumbersome, low throughput in nature, expensive and require significant volumes of biofluid, thus limiting their routine application in clinical or diagnostic settings. To address many of these limitations, flow cytometry-based strategies have been developed to allow for efficient sorting of bioparticles based on size and/or surface markers using reduced sample volumes (13). However, sorting rates are limited by the relatively low sample concentrations (necessary to avoid swarming effects) and the resolution of sEVs below 500 nm requires the adoption of fluorescent labelling strategies (14,15).

Recently, microfluidic systems have emerged as promising tools for the fast, efficient and robust focusing and isolation of micron-sized species from biofluids. Indeed, a variety of microfluidic techniques have already been shown to be effective for such purposes. Based on the manipulating forces involved, these methods can be categorized as being either active or passive in nature. Active techniques such as dielectrophoresis (DEP) (16,17), magnetophoresis (MP) (18) and acoustophoresis (AP) (19,20) rely on the application of an external force field, whereas passive techniques solely rely on the control of geometrical and fluid properties (i.e. intrinsic hydrodynamic forces), such as pinched flow fractionation (PFF) (21), deterministic lateral displacement (DLD) (22,23) and inertial microfluidics (24). Amongst passive approaches, inertial microfluidic systems have attracted most attention due to their ability to process biofluids at extremely high volumetric flow rates (∼ml/min); which is especially useful in rare event detection applications, such as the isolation of circulating tumor cells (CTCs) (25). In the case of inertial focusing, particles of a given size are transported along a microchannel and preferentially migrate to well-defined positions across the channel cross-section (24). Such inertial approaches are well suited to label-free and high-throughput manipulation of microparticles and cells. Despite the robustness of inertial focusing in Newtonian fluids, its practical implementation for the isolation of nanoscale species has been limited, primarily due to the increasing role of Brownian motion as particle size decreases. To this end, elastic forces (stemming from the rheological properties of the processing fluid) can in principle be used to provide for enhanced control over species with sub-micron dimensions.

Viscoelastic manipulation of micron-sized particles within microfluidic channels has been used to good effect in both focusing and sorting applications. In basic terms, viscoelastic fluids exhibit both viscous and elastic characteristics when undergoing deformation. Such behaviour creates a normal stress anisotropy within the fluid, which leads to transverse forces and lateral migration of particles inside the medium. Karnis *et al.* first reported viscoelasticity-induced lateral migration of non-colloidal, buoyancy-free particles towards the flow axis in macroscale pipes (26). In subsequent studies, Leal *et al.* (27) and Brunn (28) derived models for the lateral migration velocity of rigid spheres in second order fluids, noting that the ratio of the lateral and longitudinal velocities scales with the square of the blockage ratio. Importantly, such analyses suggest a need for extended channel/pipe lengths when focusing micron-sized particles to well-defined positions, and highlight the potential of viscoelastic microfluidics for the precise confinement of particles due to the accessibility of high blockage ratios within microchannels (29). Accordingly, viscoelastic microfluidic systems have been used in a variety of particle focusing, separation and flow cytometry studies (30–39). For example, D’Avino and co-workers demonstrated through both numerical simulations and experiment that small blockage ratios result in inward migration of particles towards the centerline, whereas high blockage ratios favor wall attraction (29). In addition, by implementing a non-Newtonian sheath flow, both size- (37) and shape-based (38) particle separations have been demonstrated. Most recently, CTC isolation from untreated whole blood has been demonstrated using a non-Newtonian fluid sheath flow(39). Based on the intrinsic elastic properties of viscoelastic fluids, the need for externally imposed force fields or complex channels geometries (typical in inertial systems) are eliminated, and thus viscoelastic microfluidic systems normally comprise simple straight microchannels (40–42). Indeed, Kang *et al*. showed that an extremely dilute solution of λ-DNA can be used to confine micron-diameter particles at the center of a straight microtube due to its superior elastic properties when compared to synthetic polymer solutions (40). Furthermore, sheathless shape-based (spherical and peanut shaped) and size-based (spherical) microparticle sorting has been demonstrated using viscoelastic fluids (42, 43).

Although, many of the above studies have been successful in manipulating micron-sized objects, the focusing and separation of sub-micron species is far more challenging due to the fact that the elastic force scales with particle volume. That said, Kim *et al.* recently reported for the first time the use of viscoelastic fluids to focus 500 and 200 nm particles along a straight microfluidic channel, although no significant focusing was observed for 100 nm particles (45). Related studies also demonstrated that 200 nm particles could be focused at the center of an 8 cm long cylindrical microchannel by adjusting flow rates so that elastic forces dominate Brownian forces (46). Additionally, Liu and co-workers used a double spiral channel (with a total length exceeding 6 cm) to focus and efficiently separate two sets of binary mixtures, namely 100/2000 nm polystyrene particles and λ-DNA molecules/platelets (47). Subsequently, the same group used a viscoelastic-based method incorporating a non-Newtonian sheath fluid to isolate exosomes from serum in a continuous manner (48). Despite these successes, all the aforementioned studies required the use of channel lengths on the order of a few centimeters for the focusing of sub-micron particles (29). Moreover, the processing of species with characteristic dimensions less than 100 nm is even more challenging due to the effects of Brownian forces on smaller nanoparticles, leading the use of high pressures within extended channels.

Recently, oscillatory flows have been used to eliminate the need for long microchannel lengths and successfully applied to the inertial focusing of particles down to 500 nm (49). However, the use of oscillatory flows to manipulate species below this size is still unexplored. Mutlu *et al.* have demonstrated that oscillatory inertial microfluidics can be used to focus particles in low Reynolds number regimes (one order of magnitude less than steady-flow inertial microfluidics) (49), however, the efficient focusing of species with dimensions below 100 nm would require microchannels with reduced dimensional cross-sections and channel lengths (on the order of few millimeters or microns) to allow access to high flow velocities. Although, such structures have been fabricated in rigid epoxy(42) or silicon (50) for example, system complexity and cost are unavoidably increased.

To this end, herein we present an oscillatory viscoelastic system, incorporating a sheath-free microchannel that is able to focus and separate particles and biological species with diameters ranging from a few microns to a few tens of nanometers. **Figure 1a** shows a schematic view of the microfluidic platform, which includes a pressure driven microfluidic chip coupled with an in-house electronic circuit to generate oscillatory flow. Using such an approach, objects oscillating within the microfluidic channel can be focused at specific locations based on their size (**Figure 1b**). Specifically, we initially focus and separate 10, 5, and 1 μm diameter particles within a short (4 mm long) microchannel. The same geometry is then used to separate and process RNA granules (p-bodies) from a mammalian cell lysate sample. Subsequently, oscillatory viscoelastic flows are used to perform rapid and efficient single file focusing of 500, 200, and 100 nm diameter nanoparticles along a short length high aspect ratio microchannel. Finally, and for the first time, focusing of 40 and 20 nm diameter particles is demonstrated. The combination of viscoelastic microfluidics and oscillatory flows is shown to be highly effective in overcoming Brownian motion and is widely applicable to the precise focusing of objects with sizes as small as 20 nm. Given the performance of our experimental setup (using artificial particles), we explored the potential of oscillatory viscoelastic microfluidics in manipulating biological species, and demonstrate efficient focusing of both λ-DNA and small extracellular vesicles (< 200 nm).

**Figure 1.**
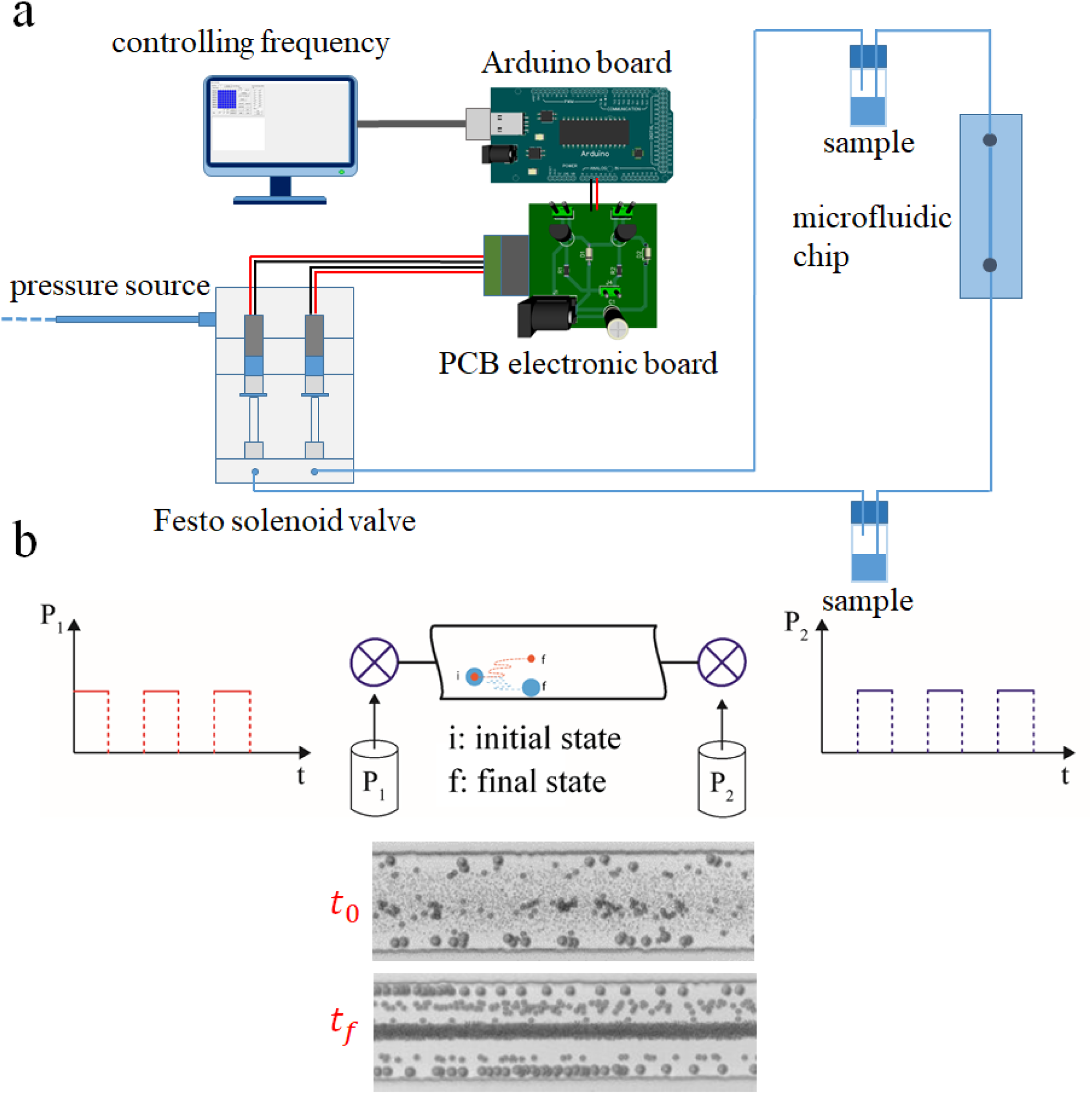
Principal of the oscillatory viscoelastic microfluidics. a) Schematic of the oscillatory viscoelastic microfluidic set-up. The system consists of solenoid valves (Festo, Zuirch, Switzerland) which are connected to a pressure source and an electronic PCB board responsible for actuating the valves to open and close states. A custom-made software is used to control the frequency and the number of oscillations. b) Oscillatory flow is generated by using the system described in part (a) and elastic forces acting on the particles cause the lateral migration and eventually focusing of them. Particles of different size (1, 5, and 10 μm) with a randomly distributed initial position (*t*_0_) can be focused at discrete parts of the microchannel after sufficient number of oscillations(*t*_*f*_).

## Results and Discussion

### Viscoelastic microparticle focusing and separation based on oscillatory flow

To examine the capabilities of oscillatory viscoelastic flows for separating micron-sized species, a ternary mixture of particles, with average diameters of 10 μm, 5 μm and 1 μm, was introduced in the microfluidic device. The effect of pressure (controlling both elasticity and inertia effect) on particle migration was studied for pressures between 1 and 2.5 bar (in 0.5 bar steps) and an oscillation frequency of 2 Hz, with particles being allowed to reach steady state positions before final images were taken. As shown in **Figure 2a**, a pressure increase results in both an increase in inertia and elasticity (defined by the Reynolds number, *Re*, and Weissenberg number, *Wi*, respectively: **Supplementary Note 1**). Since the highest *Re* accessed is 0.08 (indicating negligible effects of inertia), the driver for the lateral migration of particles is the elasticity-induced lift force. Within an oscillation period of a few seconds, 10 μm diameter particles focus near the microchannel walls, 5 μm diameter particles between the centerline and walls, with 1 μm particles being aligned to the microchannel centerline. The reason for such a migration behavior, can be understood through consideration of the blockage ratio, *β* = *a*_*p*_/*D*_*h*_, where *a*_*p*_ reports the particle size and *D*_*h*_ is the hydraulic diameter of the microchannel. Herein, a higher blockage ratio leads to a greater off-center shift of the viscoelastic focusing positions.

**Figure 2.**
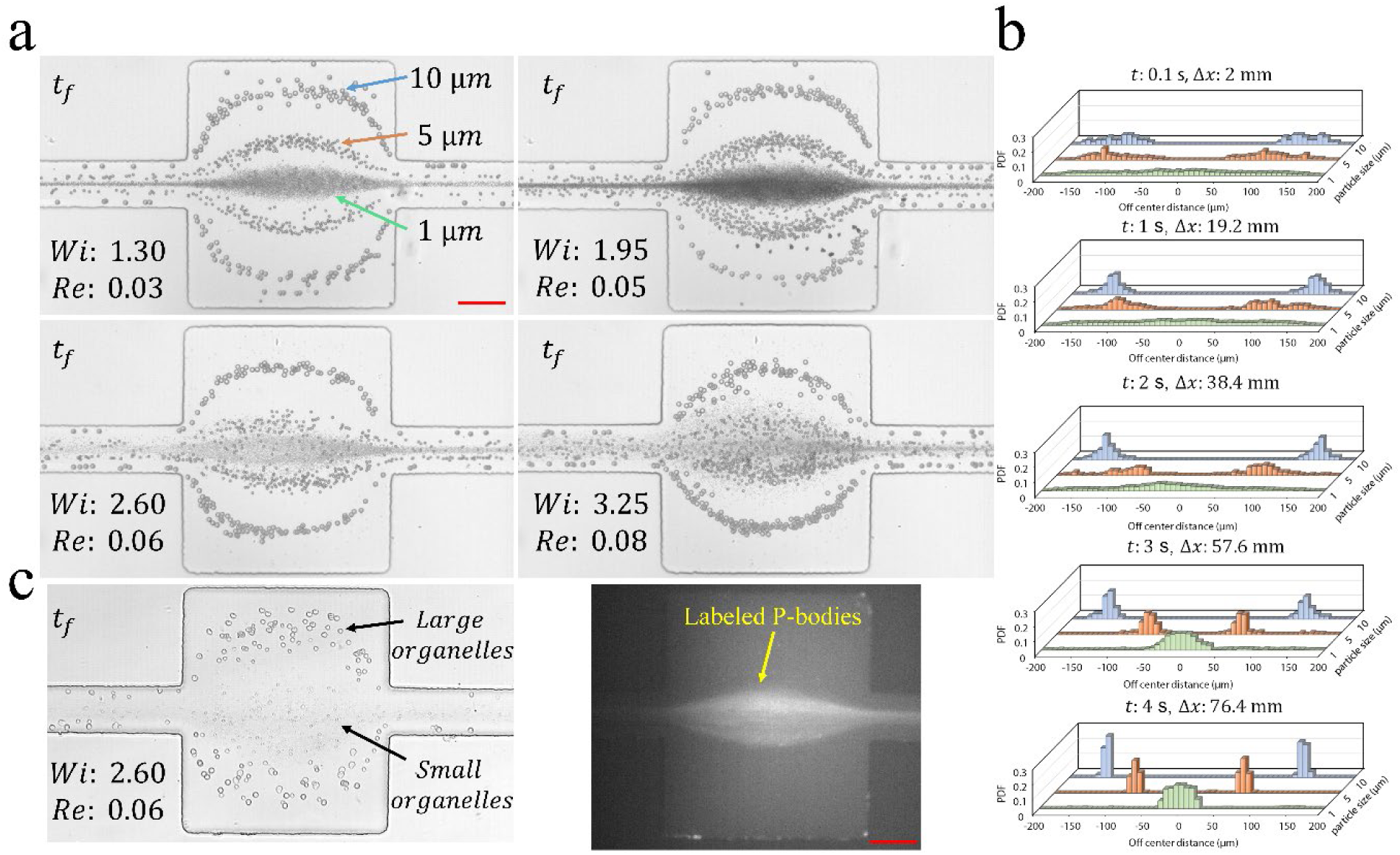
PS microparticle and bioparticle focusing and separation using oscillatory viscoelastic microfluidics. a) Effect of pressure (in terms of Wi and Re) on the focusing and separation of 1, 5, and 10 μm particles in *PEO*_400*KDa*,1%_,in a straight 80 μm-width, 12 μm-height and 4 mm length microchannel with an oscillating frequency of 2 Hz. An expansion area with 400 μm-width and 12 μm-height at the center of the microchannel is used for better visualization of the focusing and separation processes. b) Time evolution of focusing and separation of the particles in the case of Wi: 1.3 and Re: 0.03. Due to the low Re, the inertia effect can be neglected and particles migrate due to the center- and wall-directed elastic lift forces. Blockage ratio is the key component for lateral migration of the particles in different equilibrium positions. After 4 seconds of oscillation (corresponding to an equivalent channel length of 76.4 mm), high-resolution separation can be achieved. c) Cell lysate was introduced to the same microchannel as described in part (b) in a *PEO*_400*KDa*,0.5%_ solution and similar behavior was observed. Larger organelles including cell nucleus are focused near the walls, while small organelles including GFP-fluorescently labeled p-bodies are focused at the centerline. Scale bars are 80 μm.

Such behavior implies that two opposing elastic forces affect lateral migration, i.e. a centerline-directed elastic force, *F*_*el*→*c*_, and a wall-directed elastic force *F*_*el*→*w*_ (44). Competition between the two components of the elastic lift force determines the lateral focusing position of particles on the basis of their size and flow rate. Based on the results shown in **Figure 2**, smaller particles (having a low blockage ratio) tend to focus at the centerline due to the dominance of *F*_*el*→*c*_ over *F*_*el*→*w*_ (44). On the other hand, since *F*_*el*→*w*_ is more strongly dependent on the particle size than *F*_*el*→*c*_ (44), larger particles (having high blockage ratios) tend to migrate to off-center focusing positions due to stronger wall-directed elastic forces (*F*_*el*→*w*_). As shown in **Figure 2a**, the highest focusing efficiency takes place when *Wi* = 1.3 and *Re* = 0.03. An increase in pressure (*Re* = 0.03 to 0.08 and *Wi* = 1.3 to 3.25) causes larger particles to migrate toward the centerline, with smaller particles diverging from it.

**Figure 2b** presents the time evolution of oscillatory focusing and separation of the particles for the case when *Wi* =1.3 and *Re* = 0.03 (see **Supplementary Movie S1**). All plots were extracted from the expansion part of the microchannel and acquisition commenced after 100 ms, i.e. when particles have already traveled a distance of 2 mm from the inlet. Even after such a short period of time, the 10 μm diameter particles have already started to migrate towards the microchannel walls, 5 μm particles have been depleted from the center and are accumulating in a semi-focused state, with the 1 μm particles being dispersed across the expansion zone. After 4 seconds of oscillation, all particles have reached their final equilibrium positions and have travelled an effective distance of 76,800 μm; a sufficient length to ensure complete separation. (**Supplementary Movie S2**).

As a further proof that the oscillatory focusing platform is capable of separating micron-sized species over short path lengths, we then applied this method to a complex and relevant biological sample. Specifically, we attempted to purify p-bodies, cytoplasmic RNA granules that are involved in RNA metabolism and have sizes ranging from 300 nm to 1 μm (51). For this purpose, we introduced lysate from cells expressing mNG-AGO2 (AGO2 being known to be enriched in p-bodies (52)) into the oscillatory system. The lysate contains nuclei (∼6 μm in size), debris and organelles, including p-bodies (∼1 μm in size) that are stable in the lysis solution. Since biological particles are significantly more deformable than rigid particles, a 0.5% PEO polymer solution was used (to decrease elastic effects). As shown in **Figure 2c** (Bright field image), nuclei and large organelles are concentrated near the walls, in a similar manner to large, rigid particles, since their size is comparable to the channel height, whilst p-bodies and smaller organelles are focused at the centerline of the channel. This is further demonstrated by the observation of fluorescently mNG-labelled p-bodies at the center outlet (fluorescence image in **Figure 2c**). Indeed, in a similar manner to 1 μm rigid particles, we demonstrate that p-bodies begin to focus at the center of the microchannel after 4 seconds of oscillation (**Figure S1**). In conclusion, we demonstrate for the first time, efficient focusing of cellular organelles directly from a complex biological fluid using oscillatory viscoelastic focusing

### Throughput of oscillatory viscoelastic microfluidic platform

On first inspection, a potential weakness of the oscillatory platform stems from its low throughput, since the oscillation occurs within a single channel. However, this issue can be easily mitigated by fabricating an array of parallel microchannels to realize throughputs comparable to a continuous flow focusing device. As shown in **Figure S2b**, a device incorporating 100 parallel channels (where *w* = 80 μm, *h* =12 μm and *L* = 4 mm) can be simply fabricated using photolithographic methods. Under continuous-flow operation (P=2.5 bar), focusing of 1 μm particles is not possible due to the reduced length of the microchannel. Conversely, when operating under oscillatory flow conditions (oscillation frequency of 1 Hz) with same flow condition (P=2.5 bar), particles can be focused after only 4 seconds of oscillation, with a two orders of magnitude increase in analytical throughput when compared to oscillatory flow inside a single microchannel (increase of throughput from 50 counts/s to around 5000 counts/s). Indeed, as shown in **Figure S2a**, the parallel fluidic circuit is analogous to an electric circuit, and thus an increase in the number of microchannels does not intensify the resistance of the whole system when using pressure pump as a source.

### Viscoelastic nanoparticle focusing based on oscillatory flow

Next, we used the oscillatory viscoelastic microfluidic platform to focus nanoparticles, segmenting our study into two size regimes, namely nanoparticles with dimensions equal to or larger than 100 nm (sub-micron regime) and nanoparticles with dimensions smaller than 100 nm (nano-regime). Initially, particles with diameters of 100 nm, 200 nm, and 500 nm were processed using an oscillation frequency of 2 Hz and pressures between 1 and 2.5 bar. The microfluidic device used for study of this size regime comprises a 4 mm long microchannel, having a height of 1.4 μm and a width of 20 μm. **Figure 3** summarizes nanoparticle focusing behavior under the investigated conditions. At the highest pressure, *Re* is 0.00017, indicating that inertial effects are negligible. However, *Wi* values range from 0.26 (at 1 bar) to 0.65 (at 2.5 bar) which seems to be sufficiently high to induce lateral migration and focusing of nanoparticles. Indeed, for all particle diameters, we observed complete focusing at pressures of 2 bar and 2.5 bar, as shown by the fluorescence images obtained after 40 seconds (**Figure 3a-c**). Specifically, for 100, 200 and 500 nm nanoparticles at a pressure of 2.5 bar, focusing efficiencies were 86%, 85%, and 90%, respectively. Accordingly, pressure values between 2 and 2.5 bar (0.52 < *Wi* < 0.65) allow for the efficient focusing of nanoparticles with diameters between 100 and 500 nm (and blockage ratios between 0.04 and 0.2) at the channel centerline.

**Figure 3.**
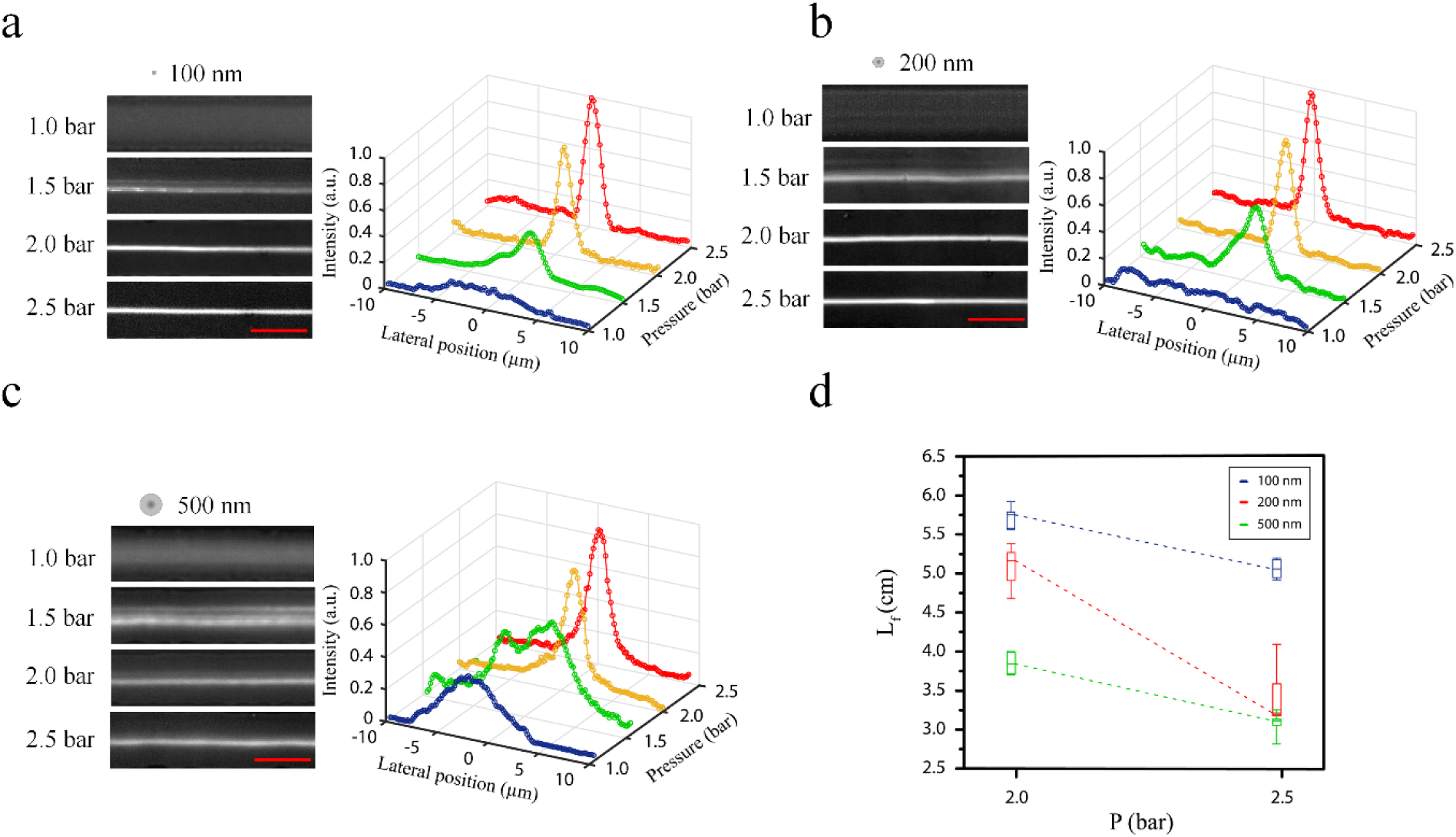
Effect of pressure on the focusing efficiency of 100, 200, and 500 nm particles within oscillatory viscoelastic microfluidics. The microchannel was fabricated using two-Photon Polymerization lithography with dimensions of 20 μm-width, 1.4 μm-height and 4 mm length. a, b, c) Effect of pressure (in terms of Wi) on the focusing efficiency of nanoparticles in *PEO*_400*KDa*,1%_. In each case, images were recorded after 40 seconds of oscillation to ensure final focusing state. The Wi numbers for pressures at 1, 1.5, 2, and 2.5 bar are 0.26, 0.39, 0.52, and 0.65 respectively. Increasing the pressure results in dominant elastic forces enabling lateral migration of nanoparticles to minimum shear rate position. Since a high aspect ratio channel is used, most of the particles tend to migrate in the centerline. Blockage ratio values for 100 nm, 200 nm, and 500 in such microchannel are 0.04, 0.08, and 0.2 respectively. d) Diagram of focusing length vs pressure for 100 nm (β: 0.04), 200 nm (β: 0.08), 500 nm (β: 0.2) nanoparticles. In the case of 2 bar (Wi: 0.52), *L*_*f*_ values for 100 nm, 200 nm, and 500 nm are 5.7 cm, 5.1 cm, and 3.8 cm, respectively. Increasing the pressure to 2.5 bar (Wi: 0.65) results in faster focusing of the particles with similar trend and *L*_*f*_ values for 100 nm, 200 nm, and 500 nm are 5.1 cm, 3.4 cm, and 3.1 cm, respectively. Scale bars are 20 μm.

We next estimated the focusing length, *L*_*f* =_ *Ut*_*f*_, where *t*_*f*_ is the time required for particles to reach a stable minimum FWHM and *U* is the longitudinal velocity of particles travelling at the centerline. As illustrated in **Figure 3d**, this yielded focusing lengths at 2 bar (*Wi* = 0.52) of 5.7 cm for 100 nm particles (*β* = 0.04), 5.1 cm for 200 nm particles (*β* = 0.08) and 3.8 cm for 500 nm particles (*β* = 0.2). At 2.5 bar (*Wi* = 0.65), focusing lengths for 100 nm, 200 nm and 500 nm particles decreased to 5.1 cm, 3.4 cm and 3.1 cm, respectively, due to the higher exerted elastic forces. This analysis illustrates the fact that as the particle size is reduced, a longer channel is needed for centerline focusing, and that an increase in pressure (or elasticity) accelerates the focusing process. Next, the time evolution of the focusing process was analyzed over a period of 40 seconds. For 100 and 200 nm particles, single file focusing was accomplished in 30 seconds, whilst 500 nm particles were focused into a single file configuration within 20 seconds (**Figure 4a-c**). **Supplementary Movies S3** and **S4** illustrate the time evolution and final focusing state of 500 nm particles. One of the most useful features of oscillatory viscoelastic microfluidic systems is the facility to perform single particle tracking. As shown in **Figure 4d**, characteristic focusing times extracted from single particle trajectories (from near the wall to the focusing position) were 30 seconds for 100 nm particles, 24 seconds for 200 nm particles and 18 seconds for 500 nm particles. An additional advantage of oscillatory viscoelastic microfluidic systems is the ability to access high flow velocities (to improve focusing efficiency) through the use of short channel lengths. For instance, in the current study a 4 mm long channel was used for all experiments. For an analogous continuous flow system incorporating the same microchannel cross-section, a length of approximately 4 cm (and a 10-fold increase in pressure) would be needed to attain an equivalent focusing efficiency.

**Figure 4.**
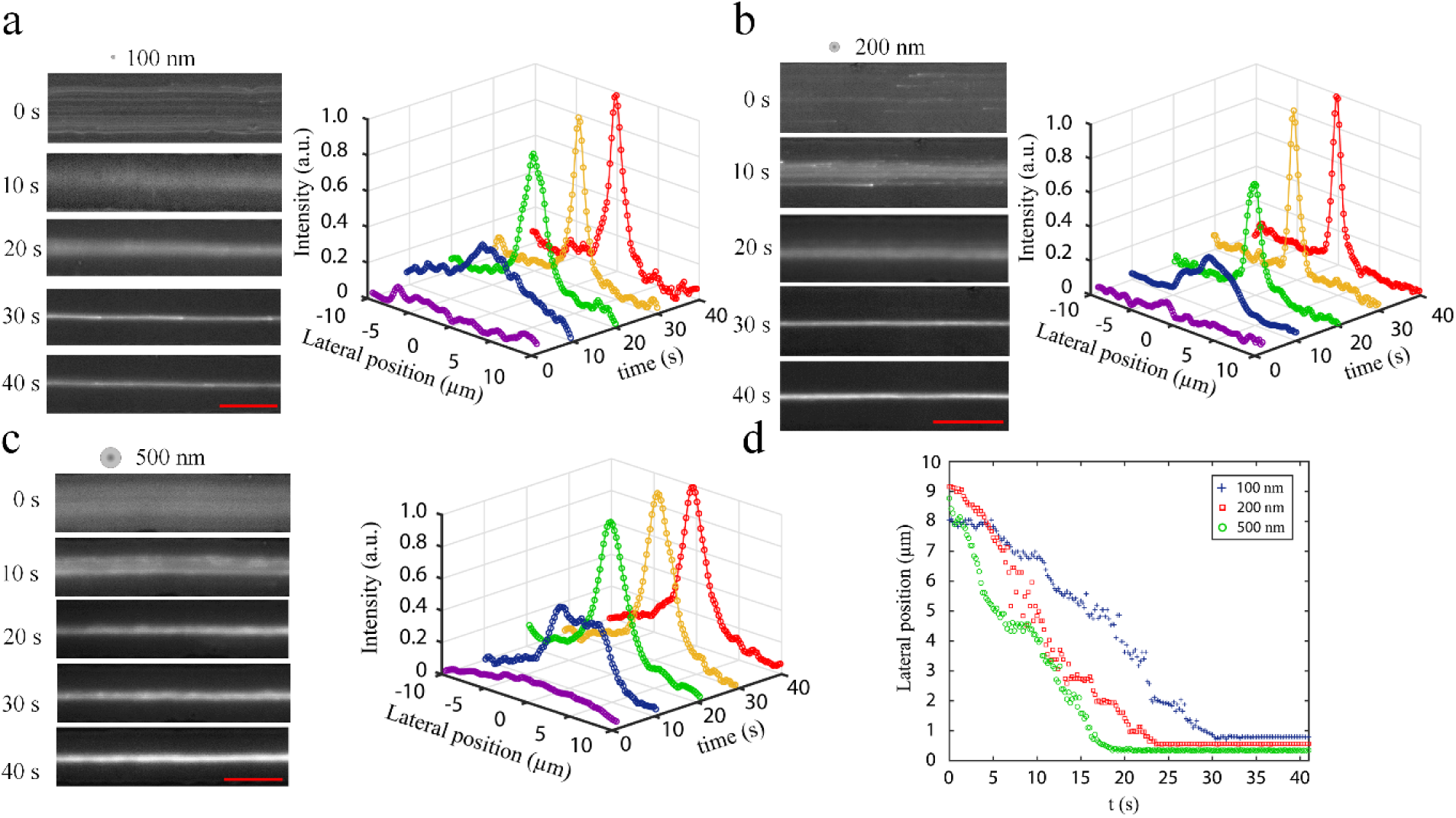
Time evolution of nanoparticles in oscillatory flow microfluidics vs focusing efficiency of a) 100 nm b) 200 nm and c) 500 nm nanoparticles. a) For 100 nm particles and pressure of 2.5 bar (Wi: 0.65), a time period of 30 seconds is adequate to reach the final focusing state. Similarly, for 200 nm (b), and 500 nm particles (c), time periods of 30 and 20 seconds are needed for the final focusing state. d) Single nanoparticle tracking reveals similar behavior. In the case of 100, 200, and 500 nm, it takes approximately 18, 24, and 30 seconds for the individual particles to migrate from the wall (around 8 μm away from center to center) to the centerline. Scale bars are 20 μm.

To further investigate the effect of the blockage ratio on particle migration within the nano-regime, the platform was used to process 200, 500 and 1000 nm diameter nanoparticles flowing through high aspect ratio microchannels. Representative fluorescence images and corresponding intensity profiles at different blockage ratios are shown in **Figure S3**. As previously noted, for a microchannel with a height of 1.4 μm, both 200 nm (β: 0.076) and 500 nm (β: 0.19) nanoparticles were focused at the center of the microchannel (**Figure S3**; left). For the larger 1μm particles, the blockage ratio is higher (β: 0.382), and thus they are aligned at off-center focusing positions regardless of their initial positions, whilst 500 nm (β: 0.19) particles are focused at the centerline (**Figure S3**; bottom). This behavior can be observed from the images recorded at the final equilibrium condition and the corresponding normalized fluorescence intensity (NFI) profiles for both 500 nm (one peak) and 1μm (two curve profile) particles. By decreasing the channel height to 700 nm and using a pressure of 4 bar, two off-center focusing positions are formed in the case of 500 nm particles (β: 0.137), while 200 nm nanoparticles (β: 0.148) are squeezed at the centerline (**Figure S3**; right). Accordingly, oscillatory viscoelastic microfluidics should allow for the efficient separation of nanometer-sized bioparticles, such as exosomes from extracellular vesicles or cell free DNA from plasma and platelets.

It is now recognized that the focusing, detection and isolation of bioparticles with dimensions below 100 nm represents one of the most important challenges in the field of cell-free RNA diagnostics (1). In this regard, the realization of a method able to robustly focus and isolate nanoparticles with hydrodynamic radii below 100 nm will potentially revolutionize our ability to isolate and process sEVs. Initial experiments aimed at focusing 20 nm and 40 nm diameter nanoparticles in microchannels having a width of 20 μm, a height of 1.4 μm and length of 4 mm, failed to demonstrate efficient focusing performance. Accordingly, the channel height was reduced to 700 nm, whilst keeping all other dimensions constant. Using such a device, the effect of pressure on focusing efficiency was first evaluated, with images taken after 40 seconds of oscillation at a frequency of 2 Hz. As can been seen in **Figure 5a**, 40 nm particles begin to align at the center of the channel at a pressure of 3.5 bar (*Wi* = 0.47, *β* = 0.03), with a further increase in pressure to 4 bar (*Wi* = 0.53, *β* = 0.03) resulting in an improved focusing efficiency of 61% (**Supplementary Movie S5**). Similar behavior is observed in the case of 20 nm diameter nanoparticles (β = 0.015), with a focusing efficiency of 51% being achieved at 4 bar (*Wi* = 0.53). As shown in **Figure 5b**, FWHM values of 4.95 μm and 6.95 μm were realized after an oscillation period of 40 seconds, for 40 nm and 20 nm particles, respectively.

**Figure 5.**
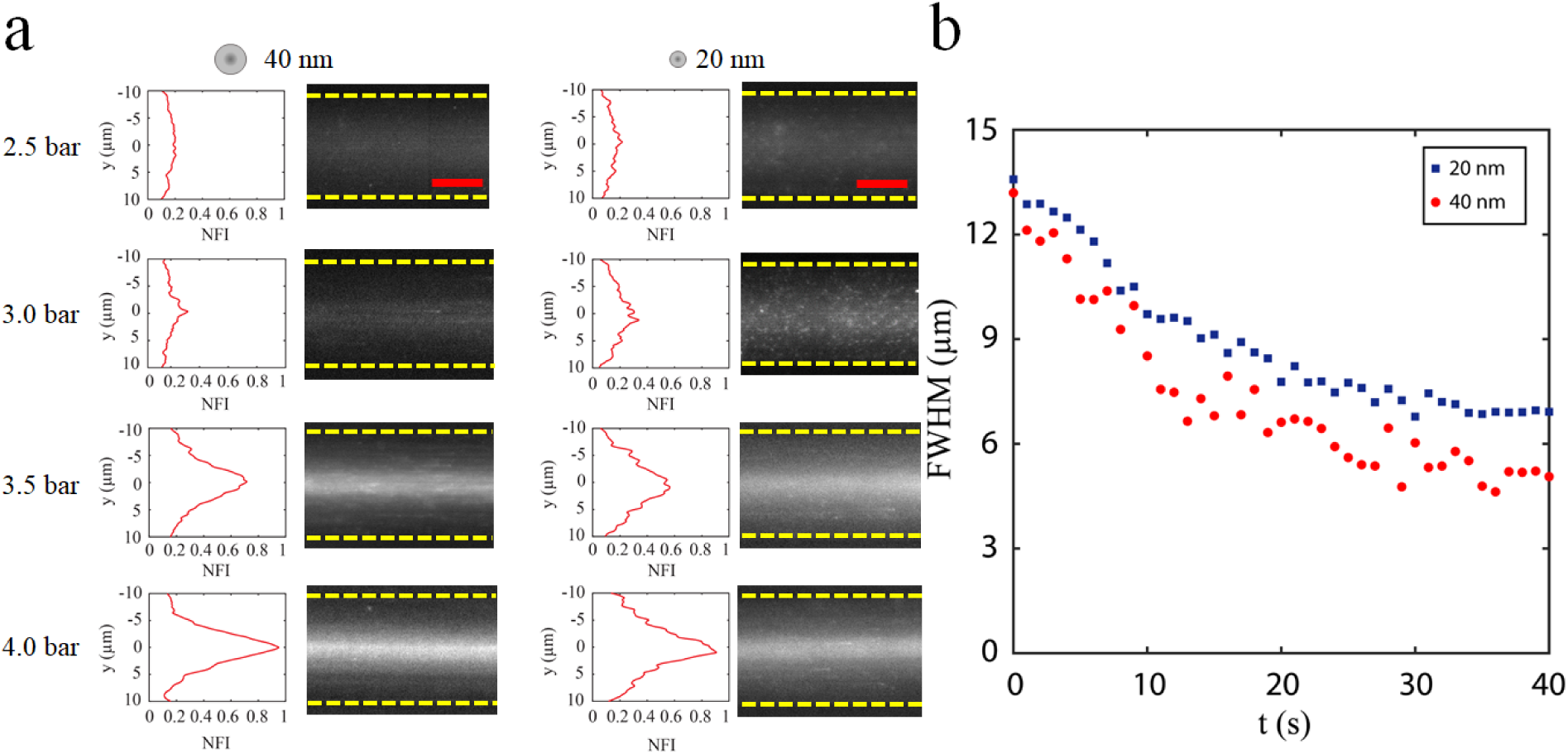
Focusing of 20 nm and 40 nm particles in oscillatory viscoelastic microfluidics. a) Effect of pressure (in terms of Wi) on the focusing efficiency of nanoparticles. For pressure values below 3.5 bar, no focusing was observed. Pressures of 3.5 (Wi: 0.47), and 4 bar (Wi: 0.53) resulted in focusing of the nanoparticles at the center of the channel. Images for each case were recorded after 40 seconds oscillation at a frequency of 2 Hz. b) FWHM values were calculated at a pressure of 4 bar case. FWHM values reached a final value of 4.95 μm and 6.95 μm for 40 nm and 20 nm, respectively. Scale bars are 10 μm.

### Viscoelastic bio-nanoparticle focusing based on oscillatory flow

Based on such excellent focusing performance, we used the oscillatory viscoelastic microfluidic platform to focus λ-DNA, and a population of mNG-labelled sEVs concentrated from cell culture conditioned medium. λ-DNA molecules have a radius of gyration of 530 nm, but are more challenging to focus at the channel centerline than rigid particles of a similar size. Experiments involving λ-DNA were performed using the sub-micron regime microfluidic device (comprising a channel that is 20 μm wide, 1.4 μm high and 4 mm long). As can be seen in **Figure S4a**, λ-DNA molecules were focused at the channel centerline (after 30 seconds). Under these conditions, *Wi*>0.52 induces sufficient elastic force to focus λ-DNA molecules. Indeed, when *Wi* = 0.65, the focusing efficiency reaches to 71.5%. Experiments involving sEVs (with a mean diameter of 122 nm; **Figure S5**) were performed using the nano-regime microfluidic device (**Figure S4b**). Using an oscillation frequency of 2 Hz and a pressure of 4 bar, a focusing time of 25 seconds and focusing efficiency of 67.4% were realized. These data confirm that biological species with dimensions ranging from a few microns to a hundred nanometers can be precisely manipulated and focused at specific regions of a microchannel through the combination of viscoelastic forces and oscillatory flows.

### Viscoelastic focusing on a diffusion-elasticity state space map

The operating flow regimes associated with nanoparticle size and viscoelastic focusing can be depicted using diffusion-elasticity state space map (**Figure 6**). Since the Reynolds number is negligible and inertial effects insufficient for lateral migration of particles, the two main forces responsible for the transverse migration of particles are the elastic force due to fluid elasticity and the Brownian force due to diffusion. These two forces can be compared using the dimensionless number, *φ*, which is defined as the ratio of lateral velocity (*u*_*p*_) due to elastic effect to the diffusion constant of the particle (*D*) within the viscoelastic carrier fluid within a given length *L*_*c*_, i.e.

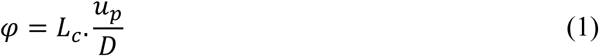

**Figure 6.**
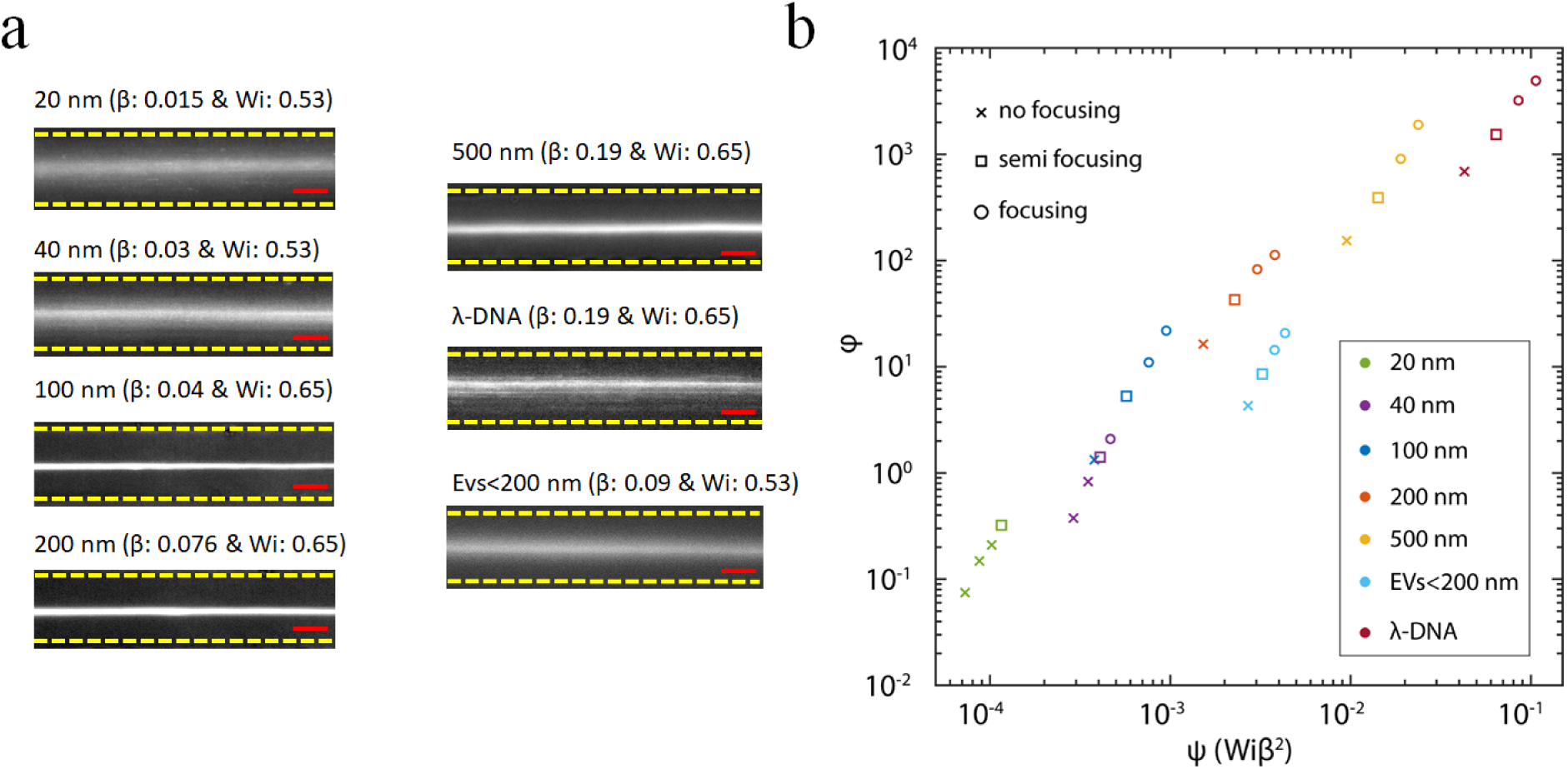
Experimental analysis of focusing phase explored in this study. a) Final focusing states at specific blockage ratios and Wi numbers for 20, 40, 100, 200, 500 nm particles, λ-DNA molecules and EVs (less than 200 nm). Scale bars are 10 μm. b) Dimensionless diffusion number (φ) versus dimensionless elastic force number (ψ) for different focusing patterns (focusing, semi focusing, and no focusing).

Ho and Leal (27) and Brunn (28, 53) have developed a theoretical approach to model particle motion in a second order fluid. By using the force equation from these models, assuming a negligible second normal stress difference (*N*_2_ ≈ 0), and considering only the first normal stress difference 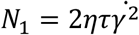, the elastic lift force on a particle can be approximated as:

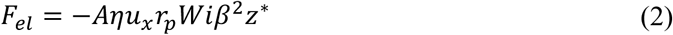

where *A* is a pre-factor, *η* the viscosity, *u*_*x*_ the mean longitudinal velocity, *r*_*p*_ radius of particle, *Wi* the Weissenberg number, *β* the blockage ratio and *z*^*^ = *z*/*H* is the dimensionless lateral position factor (54). By balancing the elastic and drag forces (*F*_*drag* =_ 6*πηr*_*p*_*u*_*p*_) and for a non-Brownian case, we can extract the lateral velocity of the particle, i.e.

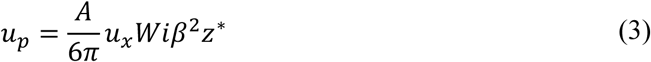

To properly describe the diffusivity of the nanoparticles in a viscoelastic fluid, short time effects (fast diffusion; see **Supplementary Note 3**) are neglected and the generalized Stokes-Einstein relation is adopted as follows (46, 55),

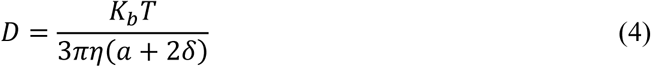

where *K*_*b*_ is the Boltzmann constant, *T* the absolute temperature, *a* diameter of the particle, δ is the thickness of depletion layer created around the nanoparticle (see **Supplementary Note 3**) within the viscoelastic fluid and *η* the macroscopic viscosity of the polymer solution. By replacing **Equation 3** and **Equation 4** in **Equation 1** and considering the characteristic length (*L*_*c*_) as the hydraulic diameter of the microchannel (*D*_*h*_), we get

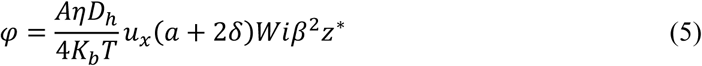

This dimensionless number (*φ*) provides a measure of the relative magnitudes of the elastic and Brownian forces. *A* is a pre-factor, and according to studies by Ho and Leal, is equal to 5*π*, whilst Brunn reports a value of 22*π*/5 (27, 53). Herein, the values of *A* and *z*^*^ are chosen to be 5*π* and 1, respectively to allow comparison of *φ* values for different particle sizes, channel dimensions, flow conditions and elasticity.

The lateral migration velocity in a viscoelastic fluid scales as *ψ* = *Wiβ*^2^ (27, 53). Accordingly, **Equation 5** can be written as:

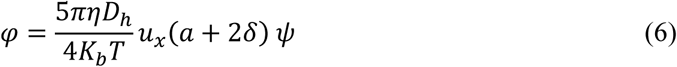

Hence, *φ* values were calculated for Brownian particles (i.e. 20, 40, 100, 200, 500 nm nanoparticles), EVs less than 200 nm in diameter and λ-DNA molecules, and a diffusion-elasticity state space map constructed (**Figure 6b**). For 20 and 40 nm particles, the magnitude of elasticity and Brownian motion are similar and diffusion is not negligible. However, an increase in *Wi* leads to a better focusing state as shown in **Figure 6b**. Interestingly, for the case of 40 nm particles we noticed that an increase in pressure initially leads to a transition between a non-focusing state and a semi-focusing state, and then a transition to focusing state (when φ≈ 2). For 20 nm particles, the focusing condition was improved to a semi-focusing state for the flow condition explored in this study (φ≈ 0.3). For the remaining nanoparticles, *φ* values are much higher than 1 even at low flow rates, suggesting efficient suppression of Brownian motion for particles in the sub-micron regime. Accordingly, particle focusing can simply be improved by increasing the pressure (or *Wi*). This leads to a considerable elastic force that induces lateral migration of particles along the centerline of the channel, and (for 40, 100, 200, 500, EVs, and λ-DNA particles) a transition from a non-focusing state to a semi-focusing state and finally to a single file focusing state. Such a simplified dimensional analysis of diffusion and elasticity for flow conditions, rheological properties and channel geometry is highly useful. However, in more complicated cases (e.g. elasto-inertial regimes or flow within curved conduits) this model should be generalized.

## Conclusion

In this study, we have demonstrated the utility of oscillatory viscoelastic microfluidic systems for manipulating and focusing objects that range in size from 20 nm up to a few microns. This study addresses three key size-regimes, namely the micron, sub-micron and nano-regime. Results show that 1, 5, and 10 μm particles could be efficiently focused at distinct parts of a (micro regime) microchannel through control of the blockage ratio. When cell lysate was introduced to the same platform, we achieved focusing of cellular organelles (p-bodies) with a similar efficiency to that observed for rigid particles. Within the sub-micron regime, 100, 200, 500 nm diameter nanoparticles, and λ-DNA molecules were successfully focused at the center of a microchannel using oscillatory viscoelastic microfluidics. Critically, and through simple modification of channel dimensions, this method could also be used for the effective separation of nanoparticles. Finally, within the nano-regime, 20 and 40 nm diameter particles, and sEVs (smaller than 200 nm in size) were also focused in a rapid and robust manner. To conclude, a dimensionless parameter (*φ*) was introduced to compare Brownian with elastic forces, and used to show that the combination of a viscoelastic fluid and oscillatory flow suppresses Brownian motion. The presented oscillatory viscoelastic microfluidic methodology enables particle manipulation over a wide size range, and has significant immediate utility in the rapid isolation of EVs, cell-free DNA, viruses, protein aggregates, λ-DNA, cellular organelles (RNA granules), and various types of cells.

## Materials and Methods

### Device Design and Fabrication

Two methods were used in the fabrication of microfluidic master molds. First, conventional lithographic techniques were used to create fluidic structures with micron-scale dimensions. Briefly, microchannel patterns were designed using AutoCAD 2018 (Autodesk, San Rafael, USA) and printed onto a 177 μm-thick fine grain emulsion film (Micro Lithography Services Ltd, Chelmsford, United Kingdom) to form a photomask. This photomask was then utilized to pattern a SU-8 coated silicon wafer (Microchem Corporation, Westborough, USA). The final master structure consisted of an inverted straight rectangular microchannel (width: 80 μm, height: 12 μm) with a total length of 4 mm and an expansion channel (width: 400 μm, height: 12 μm) to facilitate the visualization process during image acquisition. Smaller fluidic features were created using a Photonic Professional GT2 two-Photon Polymerization 3D printer (Nanoscribe GmbH, Stutensee, Germany). Specifically, the printer was used to directly form microfluidic structures on ITO coated glass using a 63× objective and IP-Dip photoresist (Nanoscribe GmbH, Stutensee, Germany). After the printing was completed, the mold was developed in poly(ethylene glycol) methacrylate (Sigma, Buchs, Switzerland) for 15 minutes and cleaned in isopropanol for a further 5 minutes. The final master structure consists of an inverted 4 mm long microchannel, having a height of 1.4 μm and a width of 20 μm. The smallest master mold consisted of an inverted 4 mm long microchannel, having a height of 700 nm and a width of 20 μm.

Subsequently, a 10:1 mixture of polydimethylsiloxane (PDMS) monomer and curing agent (Sylgard 184, Dow Corning, Midland, USA) was poured over the master-mold and peeled off after polymerization at 70°C for 4 hours. Inlet and outlet ports were created using a hole-puncher (Technical Innovations, West Palm Beach, USA). Afterwards, the structured PDMS substrate was bonded to a glass substrate (Menzel-Glaser, Monheim am Rhein, Germany) after treating both surfaces in an oxygen plasma (EMITECH K1000X, Quorum Technologies, United Kingdom) for 60 seconds. PDMS devices used for the micron and sub-micron regime experiments were bonded to a 1 mm thick glass, whilst a 200 μm thick glass substrate was used for nano-regime experiments.

### Flow Control Instrumentation

A home-built system was used to generate and control oscillatory flows within microfluidic devices (**Figure 1a**). The system comprised an UNO REV3 Arduino board (Distrelec, Uster, Switzerland), a homemade digital-to-analog conversion circuit board, solenoid valves (FESTO, Esslingen Germany) and a GUI control interface developed in Microsoft Visual Studio using C++. Commands from the GUI interface are sent to the Arduino board and used to control the solenoid valves at frequencies up to 20 Hz.

### Viscoelastic Fluid Preparation and Characterization

Viscoelastic fluids were prepared by completely dissolving polyethylene oxide (PEO, *M*_*w*_= 400kDa; Sigma-Aldrich, Buchs, Switzerland) in deionized water to a concentration of 1% (w/v). Solutions produced in this manner were then aged at 4°C for one week to realize a steady-state viscosity. Viscosities of all fluids (**Figure S6a**) were measured at room temperature using a rheometer (MCR 502, Anton Paar, Germany) and a double gap cylinder (DG 26.7). Deionized water exhibited a constant-viscosity of 0.89 cP. The viscosity of *PEO*_400*KDa*,1%_ was measured to be between 33.2 and 10.3 cP, exhibiting shear-thinning behavior. Additionally, the viscosity of *PEO*_400*KDa*,4%_ was measured to be between 846.3 and 110.1 cP, showing stronger shear-thinning behavior. Finally, the shear stress-strain curve was measured and used to uncover concentration dependent hysteresis effects. **Figure S6b** shows the shear stress-strain dependence for both of the *PEO*_400*KDa*,1%_ and *PEO*_400*KDa*,4%_ samples. It can be seen that hysteresis is significantly smaller for the *PEO*_400*KDa*,1%_ sample. High hysteresis can result in asymmetric and unpredictable lateral motion of the particles during oscillation. Considering both low hysteresis and low shear-thinning for *PEO*_400*KDa*,1%_, we chose *PEO*_400*KDa*,1%_ as the viscoelastic carrier fluid.

### Working Principle and Data Acquisition

Samples were loaded into glass vials (ThermoFisher Scientific, Waltham, USA) and delivered to the microfluidic device at a given pressure using an air pressure source. The microfluidic device was mounted on an inverted Eclipse Ti-E microscope (Nikon, Zürich, Switzerland) equipped with an IDT high speed camera (Motion Pro Y5.1, Niederoenz, Switzerland) and an ORCA-flash 4.0 CMOS camera (Hamamatsu, Solothurn, Switzerland). Bright-field imaging of particle populations and cell lysate was carried out using the high speed camera, plasma light source illumination (HPLS200 series Thorlabs, New Jersey, USA) and a 20×, 0.45 NA S Plan Fluor objective (Nikon, Zürich, Switzerland). Fluorescence imaging was performed using two different objectives. For the visualization of 100 nm, 200 nm, 500 nm particles, p-bodies and λ-DNA, a 20× 0.45 NA objective lens was used. In addition, a 60× 1.2 NA WI Plan Apo VC objective (Nikon, Zürich, Switzerland) was used to image 20 nm and 40 nm particles and extracellular vesicles. Images were processed with ImageJ software (U. S. National Institutes of Health, Bethesda, USA) using the “z-projection” and “sum slices” options. To assess focusing performance, full width at half maximum (FWHM) values were extracted from intensity profiles, with focusing efficiency being defined as the fraction of species within 1/4 microchannel width from the centerline. For bright-field experiments, the probability distribution function (PDF) was obtained by measuring fraction of the particles in the expansion region.

### Sample Preparation

Polystyrene (PS) beads with an average diameter of 10, 5, and 1 μm (Sigma-Aldrich, Buchs, Switzerland) as well as fluorescent particles with average diameters of 500, 200, 100, 40, and 20 nm (Invitrogen, Carlsbad, USA) were dispersed in deionized water at a concentration of 10^6^ particles/ml. For fluorescence labeling of λ-DNA (Sigma-Aldrich, Buchs, Switzerland), 0.2 μM YOYO-1 (ThermoFisher Scientific, Waltham, USA) was dissolved in an aqueous solution containing 60 μM Netropsin (Sigma-Aldrich, Buchs, Switzerland), 10 μM TBE buffer (Medicago, Uppsala, Sweden) and 2-Mercaptoethanol (Sigma-Aldrich, Buchs, Switzerland). This was subsequently added to the *PEO*_400*KDa*_ solution to a final concentration of 0.5% (w/v).

### Cell lines

HEK293T flp-in TREX cells (Life Technologies, Zug, Switzerland) were maintained at 37 °C within a (5% CO2) Dulbecco’s Modified Eagle Medium (Life Technologies, Zug, Switzerland) containing 4500 g/mL glucose, and further supplemented with 10% fetal bovine serum (FBS) (Life Technologies, Zug, Switzerland), 100 U/mL of penicillin (Sigma-Aldrich, Buchs, Switzerland) and 100 μg/mL of streptomycin (Sigma-Aldrich, Buchs, Switzerland).

### Cell lysis for p-body purification

First, 8×10^7^ HEK293T Flp-in TREX cells, stably expressing the mNeonGreen (mNG)-tagged Ago2 transgene, were seeded in 15-cm dishes with 2 ug/mL doxycycline (DOX) (Sigma-Aldrich, Buchs, Switzerland) being used to induce protein expression. Two days after seeding, approximately 3×10^7^ million cells were trypsinized and washed twice with PBS buffer before snap freezing the cell pellet, and subsequent storage at −80 °C. Before delivery to the microfluidic device, frozen cell pellets were homogenized and lysed for 20 minutes on a rotating wheel at 4°C using the following lysis buffer: 0.15 M NaCl (Carl Roth, Arlesheimm, Switzerland), 50 mM Tris pH = 7.5 (Carl Roth, Arlesheimm, Switzerland), 15 mM MgCl_2_ (Carl Roth, Arlesheimm, Switzerland), 200 μM phenylmethylsulfonyl fluoride (Sigma-Aldrich, Buchs, Switzerland), 5% glycerol (Brunschwig, Basel, Switzerland), 0.2% Triton-X (Carl Roth, Arlesheimm, Switzerland), supplemented by one tablet of Complete Mini EDTA-free Protease Inhibitor Cocktail (Sigma-Aldrich, Buchs, Switzerland) per 7 ml of lysis buffer.

### EV purification

For EV purification experiments, HEK293T cells expressing mNG-HRAS (FSN-HRAS) or mNG-KRAS (FSN-KRAS) were conditioned to grow in Pro293a-chemically defined medium (CDM; Lonza, Visp, Switzerland) supplemented with 1% FBS medium, by reducing the FBS content by splitting (starting from 10%). Conditioned cells were split with Accutase (STEMCELL Technologies, Köln, Germany) instead of Trypsin. Pro293a-CDM was always supplemented with 100 U/mL of penicillin and 100 μg/mL of streptomycin.

To prepare conditioned culture medium (CCM) containing fluorescent microvesicles, 20×10^7^ HEK293T FT FSN-HRAS and FSN-KRAS cells (grown in 1% FBS Pro293a-CDM) were seeded in 15-cm dishes with Pro293a-CDM without FBS. The next day protein expression was induced by addition of 2 ug/mL DOX to the medium. After three days of cell growth, CCM from two 15-cm plates was collected in one 50 mL Falcon tube. To remove residual cells, CCM was spun for 5 minutes at 500 g. Supernatant was then transferred to a fresh tube, and samples were spun for an additional 5 minutes at 1000 g. Supernatant was serially filtered using syringe-filters with cut-offs of 0.8 μm and 0.2 μm, before concentrating the CCM down to 500 μL using a Centricon 70 plus centrifugal filter (Sigma-Aldrich, Buchs, Switzerland) according to the manufacturer’s protocol (concentration step: 35 minutes at 2200 g at 18°C; collection step: for 2 minutes at 1000 g at 18°C).

## Acknowledgements

The authors would like to acknowledge support from the Swiss National Science Foundation (Nr. 205321/176011/1) and ETH Zürich. We also thank Dr Thomas Schweizer and Dr Mara Saenz de Juano for assistance with rheology and TRPS measurements.

## Author contributions

M.A, S.S and A.dM conceived the project and devised the research plan. M.A developed the instrumental platform, methodology and performed all experiments. X.C fabricated all microfluidic devices. B.M and D.V.L developed the protocols for cell culturing and extracellular vesicle purification. MA, S.S, B.M and A.dM wrote the manuscript.

