## Supplementary Information for "Oscillatory Viscoelastic Microfluidics for Efficient Focusing and Separation of Nanoscale Species"

### Note 1: Theoretical background of viscoelastic microfluidics

Particles flowing in a viscoelastic fluid are subjected to an elastic lift ($F_{el}$) force due to a normal stress difference anisotropy, where the first normal stress difference ($N_1$) produces extra tension along the particle streamline and the second normal stress difference ($N_2$) creates a secondary flow over the channel cross-section. By assuming that $N_2$is much smaller than $N_1$, $F_{el}$ is proportional to variation of $N_1$ and the size of the particle, i.e. (1)

$$F_{el} \sim a^3 \nabla N_1 \sim \lambda (\frac{ah^3}{12\eta L})^3 p^3 \quad (1)$$

where $a$ is the particle diameter, $\lambda$ is the effective relaxation time (see estimation of relaxation time in Note 2), $h$ is the channel height, $\eta$ is the fluid viscosity, $L$ is channel length, and $p$ is the pressure. The channel Reynolds number (*Re*) is defined as ratio of inertial forces to viscous forces and given by,

$$Re = \frac{\rho U D_h}{\eta_0} = \frac{2\rho Q}{\eta_0 (w+h)} \quad (2)$$

Here $\rho$ is the fluid density, $U$ is the fluid velocity, $D_h$ and is the hydraulic diameter($D_h = 2wh/h + w$) of the microchannel with (*w*) as width and (*h*) as height.

According to the Hagen-Poiseuille equation, the pressure difference in a microchannel is proportional to the average flow rate and the fluidic resistance of the channel. For a rectangular microchannel with high aspect ratio, the relation between pressure difference and the flow rate is given as (2)

$$R_h = \frac{\Delta P}{Q} = \frac{12\eta L}{wh^3} \quad (3)$$

where, $\Delta P$, Q, $\eta$, $L$, $w$, and $h$ are the pressure difference between the two ends of the microchannel, the volumetric flow rate, the dynamic viscosity of the fluid, the total length of the microchannel, the microchannel width, and the microchannel height, respectively. Combination of Equation 2 and Equation 3, allows calculation of the Reynolds number for all the pressure values used in our experiment.

The Weissenberg number (*Wi*) is defined as the ratio of elastic forces to viscous forces, i.e.

$$Wi = \lambda \dot{\gamma} = \lambda \frac{2U}{w} = \frac{2\lambda Q}{w^2 h} \quad (4)$$

where $\dot{\gamma}$ is the average fluid shear rate in the width direction of the microchannel and $\lambda$ is the effective relaxation time. Implementation of all values from experiment, allows $Wi$ to be directly calculated.

### Note 2: Estimation of relaxation time

Zimm-Rouse polymer theory has been used to estimate the relaxation time. The overlap concentration of the polymer solution is estimated as $c^* = 3M_w/4\pi N_A R_g^3$, where $M_w$ is the molecular weight of the polymer, $N_A$is the Avogadro number and $R_g$is the radius of gyration of the polymer [1]. The radius of gyration for PEO-water solutions can be estimated from the empirical equation $R_g = 0.0215 M_w^{0.583}$, and is equal to 50.2 nm [2]. Accordingly, $c^*$ for $PEO_{400kDa,1\%}$ is 0.18% w/v. Previously, several studies have demonstrated that the polymer concentration in a semi-dilute solution regime is around one order of magnitude higher than the overlap concentration [3]. Therefore, we can assume that the solution used in the current experiments is within the semi-dilute regime. Within this regime, the relaxation time can be estimated by the Rouse relaxation time [4] as shown below:

$$\lambda = \frac{6\eta_0 M_w}{c\pi^2 N_A KT} \quad (5)$$

Using Equation 5, the relaxation time can be estimated to be 0.56 ms.

**Note 3: Diffusion of nanoparticles inside a viscoelastic medium**

Previous studies have demonstrated that for a particle within a polymer solution, polymer chains close to the particle surface can form a depletion layer [5]. The thickness of the depletion layer ($\delta$) can be estimated as $\delta$=3$(R^2\xi^2/R^2 + \xi^2)^{0.5}$ where $R$ is radius of the particle and $\xi$ is the blob size [5,6]. For semi-dilute PEO solutions, the blob size can be estimated as $\xi$=$R_g(^{c}/_{c^*})^{-0.75}$=14.3 nm [5]. Accordingly, depletion layer thicknesses for different nanoparticles can be extracted, and are summarized in **Table S1**.

In this analysis, a particle may be considered to be trapped inside a polymer "cave", with both particle and cave undergo Brownian motion. Ziebacz *et al.* have characterized the motion of the particle and the cave using two diffusion coefficients, namely fast diffusion ($D_0$) for the motion of the particle inside the cave and slow diffusion ($D$) for the motion of the cave with the particle enclosed in it [5]. That said, it should be noted that diffusion inside the cave does not affect the motion of the cave [5], with the particle and the cave diffusing independently [7]. As a result, diffusion of the cave is used for further calculation, with the cave size $(a + 2\delta)$ being used in diffusion equation (Equation 4 in the main text) instead of particle size $(a)$.

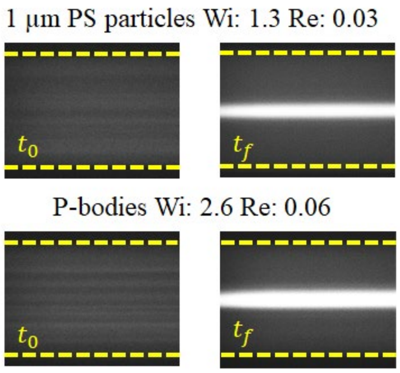


**Figure S1. Focusing of micron-sized PS and bioparticles**. Fluorescence images of 1 μm PS particles and p-bodies at $t_o$ and $t_f$.

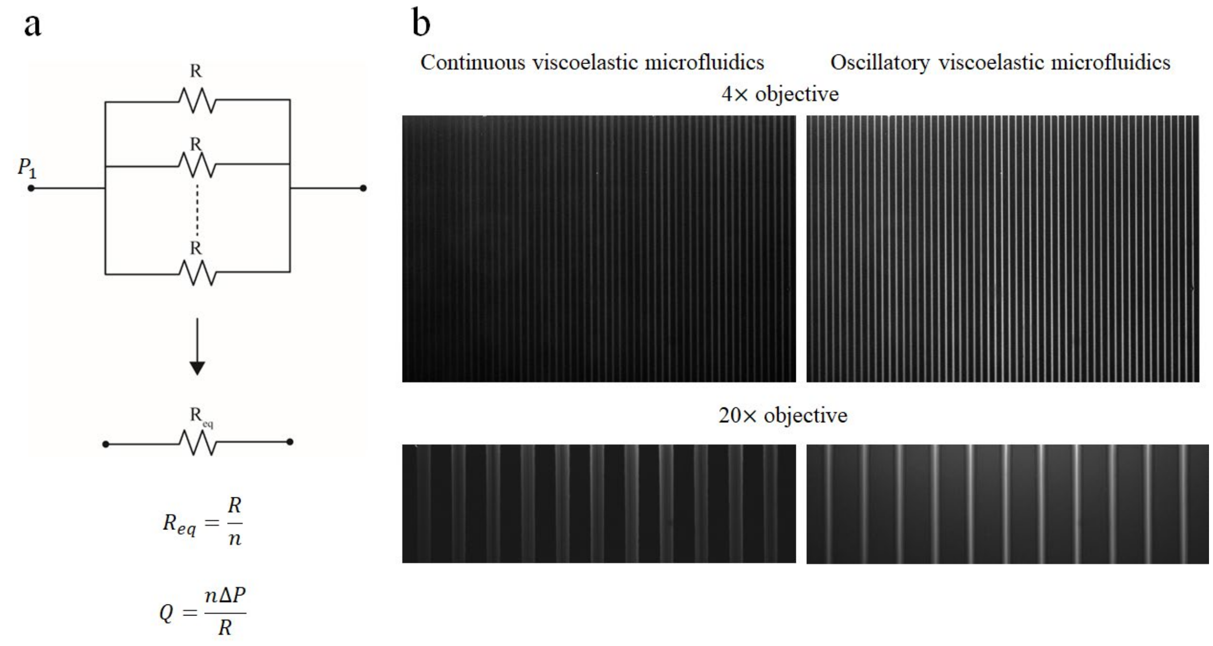


**Figure S2. Parallelization of an oscillatory viscoelastic microfluidic system**. (a) In analogy to an electronic circuit, the addition of parallel microchannel does not increase the resistance of the whole system. (b) Images of a device incorporating 100 parallel microchannels ($w$ = 80 µm, $h$ =12 µm and $L$ = 4 mm). Under continuous flow operation, no focusing of 1 µm particles was observed. However, by oscillating the flow at a frequency of 1 Hz, with an oscillation period of 4 seconds and a pressure of 2.5 bar, efficient particle alignment may be realized.

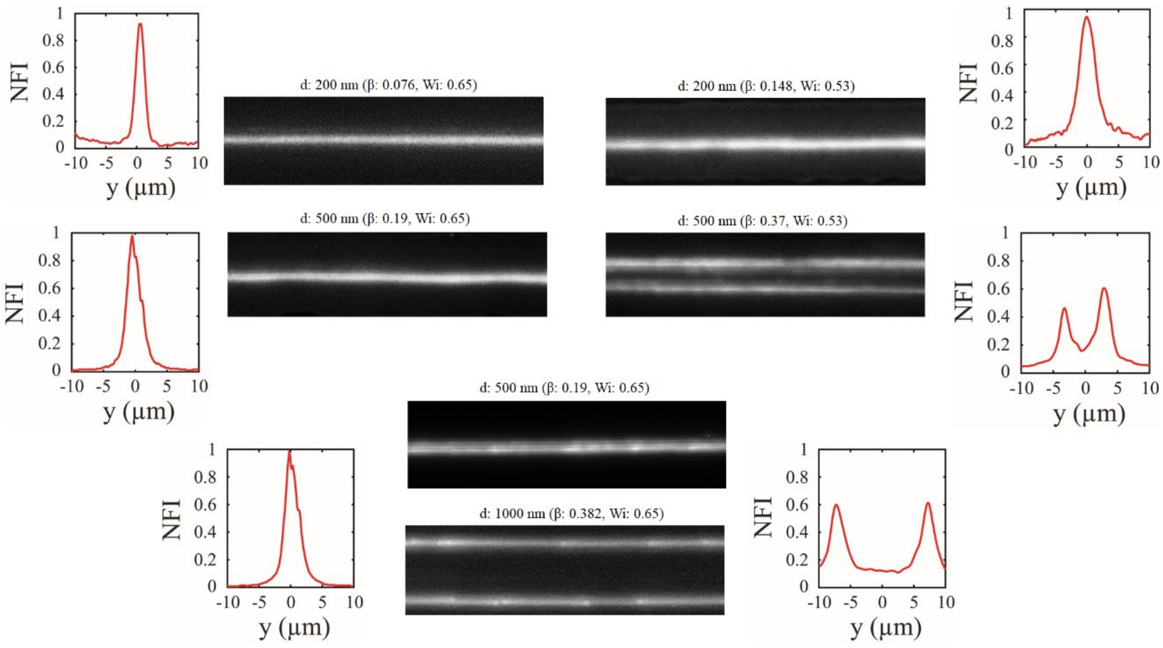


**Figure S3. Effect of channel height and blockage ratio on the focusing of nanoparticles.** For a microchannel with a height of 1.4 μm and width of 20 μm, both 200 nm and 500 nm particles are focused at the center of the microchannel (top left). For the larger 1000 nm particles inside the same microchannel, two focusing lines near the walls are observed (bottom). Inside a channel with a height of 0.7 μm and width of 20 μm, 200 nm particles are focused at the center, however, 500 nm particles start to align in two lines away from the centerline (top right).

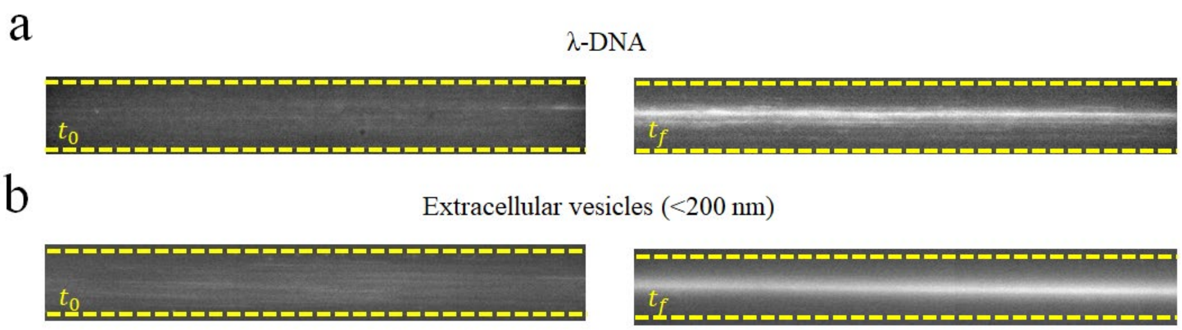


**Figure S4. Focusing of biological species**. a) Fluorescence images showing the initial and final focusing states for λ-DNA. (b) Fluorescence images showing the initial and final focusing states for sEVs.

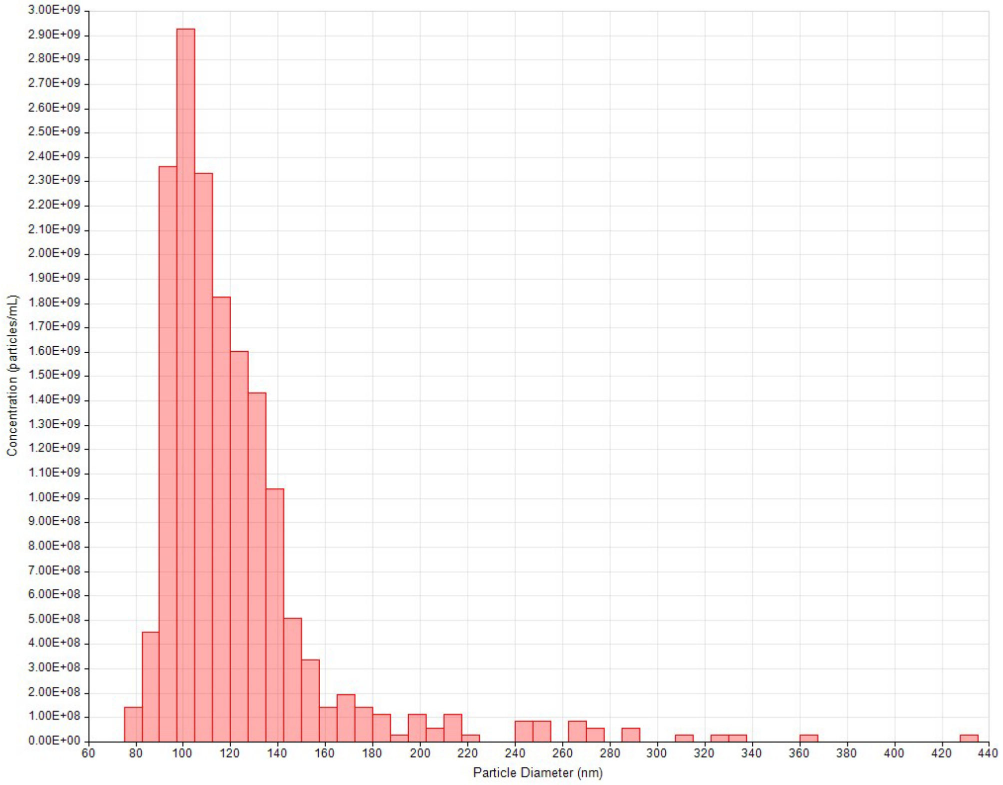


**Figure S5. Characterization of extracellular vesicle**. Vesicle size distribution plotted as a histogram of concentration versus particle diameter, obtained using a qNano Gold tunable resistive pulse sensing instrument (iZON Science, Oxford, United Kingdom). Vesicles have a mean diameter of 122 nm.

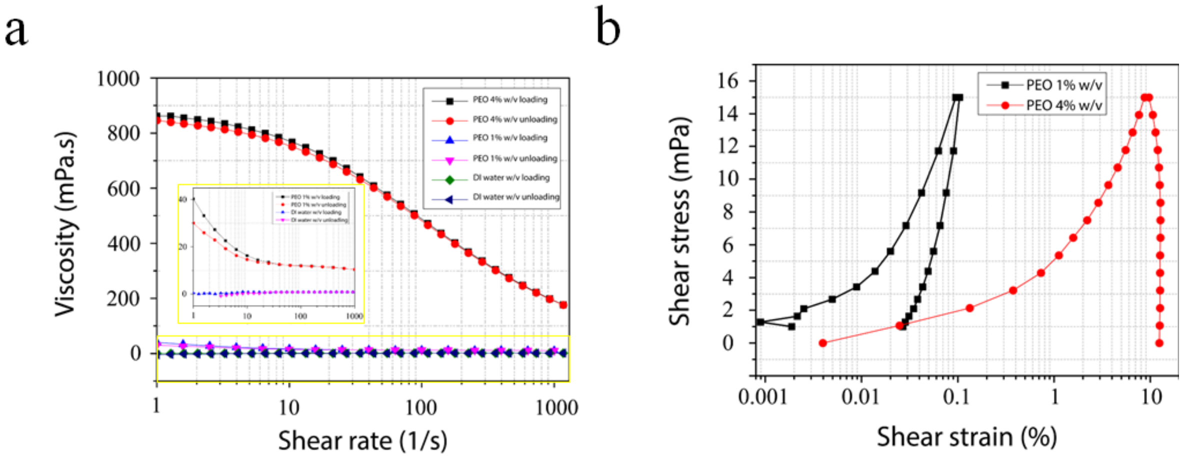


**Figure S6. Measurement of sample rheology**. (a) Viscosity as a function of shear rate for $PEO_{400kDa,1\%}$, $PEO_{400kDa,4\%}$ and DI water. $PEO_{400kDa,1\%}$exhibits minor shear thinning behavior, $PEO_{400kDa,1\%}$ behaves as a strong shear thinning medium and DI water has a constant viscosity. (b) Shear stress-strain measurements of the $PEO_{400kDa,1\%}$and $PEO_{400kDa,4\%}$ highlight weak hysteresis for $PEO_{400kDa,1\%}$and strong hysteresis for $PEO_{400kDa,4\%}$.

**Table S1**. The size of depletion layer around nanoparticles

| Particle diameter (nm) | δ (nm) |
|---|---|
| 20 | 24.6 |
| 40 | 34.9 |
| 100 | 41.3 |
| 200 | 42.5 |
| 500 | 42.9 |

**Table S2.** Dimensionless values to compare elasticity and diffusion

| 20 nm (β=0.15) | | | 40 nm (β=0.03) | | | 100 nm (β=0.04) | | | 200 nm (β=0.08) | | | 500 nm (β=0.19) | | |
|---|---|---|---|---|---|---|---|---|---|---|---|---|---|---|
| Wi | ψ | φ | Wi | ψ | φ | Wi | ψ | φ | Wi | ψ | φ | Wi | ψ | φ |
| 0.33 | $7.3\times10^{-5}$ | 0.1 | 0.33 | $2.9\times10^{-4}$ | 0.37 | 0.26 | $3.8\times10^{-4}$ | 1.3 | 0.26 | $1.5\times10^{-3}$ | 16.2 | 0.26 | $9.5\times10^{-3}$ | 152.9 |
| 0.40 | $8.7\times10^{-5}$ | 0.15 | 0.40 | $3.5\times10^{-4}$ | 0.82 | 0.39 | $5.7\times10^{-4}$ | 5.3 | 0.39 | $2.3\times10^{-3}$ | 42.8 | 0.39 | $1.4\times10^{-2}$ | 391.2 |
| 0.47 | $1.0\times10^{-5}$ | 0.21 | 0.47 | $4.1\times10^{-4}$ | 1.40 | 0.52 | $7.9\times10^{-4}$ | 10.9 | 0.52 | $3.0\times10^{-3}$ | 82.5 | 0.52 | $1.9\times10^{-2}$ | 904.7 |
| 0.53 | $1.2\times10^{-5}$ | 0.32 | 0.53 | $4.7\times10^{-4}$ | 2.08 | 0.65 | $9.5\times10^{-4}$ | 21.8 | 0.65 | $3.8\times10^{-3}$ | 112.8 | 0.65 | $2.4\times10^{-2}$ | 1898.7 |
